# Predicting circRNA-RBP interaction sites using a codon-based encoding and hybrid deep neural networks

**DOI:** 10.1101/499012

**Authors:** Kaiming Zhang, Xiaoyong Pan, Yang Yang, Hong-Bin Shen

## Abstract

Circular RNAs (circRNAs), with their crucial roles in gene regulation and disease development, have become a rising star in the RNA world. A lot of previous wet-lab studies focused on the interaction mechanisms between circRNAs and RNA-binding proteins (RBPs), as the knowledge of circRNA-RBP association is very important for understanding functions of circRNAs. Recently, the abundant CLIP-Seq experimental data has made the large-scale identification and analysis of circRNA-RBP interactions possible, while no computational tool based on machine learning has been developed yet.

We present a new deep learning-based method, CRIP (<u>C</u>irc<u>R</u>NAs <u>I</u>nteract with <u>P</u>roteins), for the prediction of RBP binding sites on circRNAs, using only the RNA sequences. In order to fully exploit the sequence information, we propose a stacked codon-based encoding scheme and a hybrid deep learning architecture, in which a convolutional neural network (CNN) learns high-level abstract features and a recurrent neural network (RNN) learns long dependency in the sequences. We construct 37 datasets including sequence fragments of binding sites on circRNAs, and each set corresponds to one RBP. The experimental results show that the new encoding scheme is superior to the existing feature representation methods for RNA sequences, and the hybrid network outperforms conventional classifiers by a large margin, where both the CNN and RNN components contribute to the performance improvement. To the best of our knowledge, CRIP is the first machine learning-based tool specialized in the prediction of circRNA-RBP interactions, which is expected to play an important role for large-scale function analysis of circRNAs.

## 1. Introduction

Circular RNAs (circRNAs) are a special type of non-coding RNAs, whose structures are characterized by non-linear backsplicing. Although categorized as noncoding RNAs, their potential to code for proteins has been reported recently [27].

Compared to linear RNA molecules, circRNAs are more stable and conserved across species [18]. Natural circRNAs have been discovered 20 years ago, while their important roles in gene regulation and disease development have attracted public attention only until recent years [17][23].

Benefiting from the high-throughput sequencing techniques, a large number of circRNA loci have been discovered in human genomes. Various databases and computational methods have been designed for circRNAs. For instance, circBase collects datasets of circRNAs and provides visualized tools for browsing circRNAs at genome scale and identifying circRNAs in sequencing data [15]; CIRCpedia also allows users to search, browse and download circRNAs with expression characteristics/features in various cell types/tissues including disease samples [37]; CircR2Disease focuses on the associations between circRNAs and diseases [13]; and CircInteractome houses the RBP- and miRNA-binding sites in human circRNA sequences [10].

According to previous studies, the regulatory functions of circRNAs largely rely on their interactions with microRNAs and RNA-binding-proteins (RBPs), acting as miRNA sponges [25][16][17] and RBP sponges [9][38]. In order to detect the interactions between proteins and RNAs, high-throughput techniques have been developed, including both *in vivo* and *in vitro* experiments [36,32]. In [24], the authors applied a soft-clustering method, RBPgroup, to various CLIP-Seq datasets, and grouped RBPs that specifically bind to the same RNA sites. [20] reported an approach circScan to identify regulatory interactions between circRNAs and RBPs by discovering back-splicing reads from cross-linking and immunoprecipitation followed by CLIP-Seq data.

Due to the availability of abundant RNA sequences, the prediction of protein-RNA interaction based on machine learning methods has been a hot topic in the bioinformatics field [21], as the data-driven methods can save the costly and laborious work of biological experiments. The prediction of protein-RNA binding sites is essentially a classification problem, involving both sequence feature representation for RNAs and classification models. The existing feature representation methods of RNAs fall into two major categories, based on statistical properties and sequence encoding schemes, respectively.

Traditionally, RNA sequence classification adopts handcrafted features, which are mainly extracted from statistical properties. For instance, *k*-tuple nucleotide composition [39] is the most basic method, which lays the foundation for a series of statistical feature extraction methods of RNAs. Note that RNAs have 4 different nucleotides, ‘A (Adenine)’, ‘G (Guanine)’, ‘C (Cytosine)’ and ‘U (Uracil)’, thus *k*-tuples have 4*^k^* different combinations, which means that each RNA sequence corresponds to a 4*^k^*-dimensional feature vector. This type of features can capture the short-range or local sequence order information [5].

During the past decade, with the rise of deep learning, sequence encoding methods have attracted more and more research attention. One-hot encoding is a simple and common feature representation method, which has been widely used in biological sequence classification [3]. For RNA/DNA sequences, each nucleotide is encoded as a 4-dimensional binary vector, which can work with both traditional classifiers and deep models. Furthermore, some researchers incorporated the secondary structure of RNAs and encoded them in the one-hot manner [31].

Besides the feature representation, various machine learning methods have been applied in the prediction of molecular interactions. For instance, support vector machines (SVMs) and random forests (RFs) have been applied to protein-protein prediction [33] and RNA-protein prediction [26]. Deep learning models have also merged, e.g., DeepBind based on CNN [1], iDeep based on multiple feature fusion [29] and iDeepE based on local and global CNNs [30].

Despite the progress on predicting interactions between linear RNAs and RBPs, the computational tools for identifying the interactions between circRNAs and RBPs have not been reported yet. Although the existing methods for linear RNAs could be applied, customized tools for circRNAs are needed due to the following reasons. First, the mechanisms of circRNAs interacting with RBPs are different from those of other types of RNAs, thus the existing methods may not be generalized well to circRNAs. Second, circRNAs have limited information for the prediction. For linear RNAs, besides the sequences, secondary structures information is usually extracted and incorporated into the predictor. Unlike linear RNAs, which have free ends and diversified secondary structure elements, circRNAs are more topologically constrained (a covalently closed continuous loop). Third, there is still room to improve the current predictors for RNA-protein interactions. For one thing, the conventional one-hot representation may lose much information of sequence patterns due to the low dimensionality and simple encoding scheme. For another, the capabilities of deep learning models are not yet fully exploited.

In this study, we propose a deep learning-based model for predicting RBP binding sites on circRNAs using sequence information alone, named CRIP, which is driven by a new stacked codon-based encoding scheme and a hybrid neural network model as the classifier. We assess the new method on the benchmark data sets of protein-circRNA binding sites, CRIP achieves higher prediction accuracy compared with both the traditional classifiers and state-of-the-art deep learning models. As far as we know, this is the first predictor for circRNA-protein interactions using machine learning, which will assist in revealing important roles of circRNAs in gene regulation.

## 2. Methods

In this study, we design a hybrid neural network model to predict the interactions between circRNAs and proteins. We first collected RBP binding sites in circRNA sequences from the CLIP-Seq experimental data. Instead of using the traditional one-hot encoding method, we propose a stacked codon-based encoding to get an initial representation for the RNA sequences. Then we use the CNN to learn high-level features from the initial representation and the long short-term memory (LSTM) network to learn dependency within the sequences. Finally, two fully connected layers are used to determine whether the given circRNA fragment is a binding site or not. The whole pipeline is shown in Fig. 1.

**Fig. 1:**
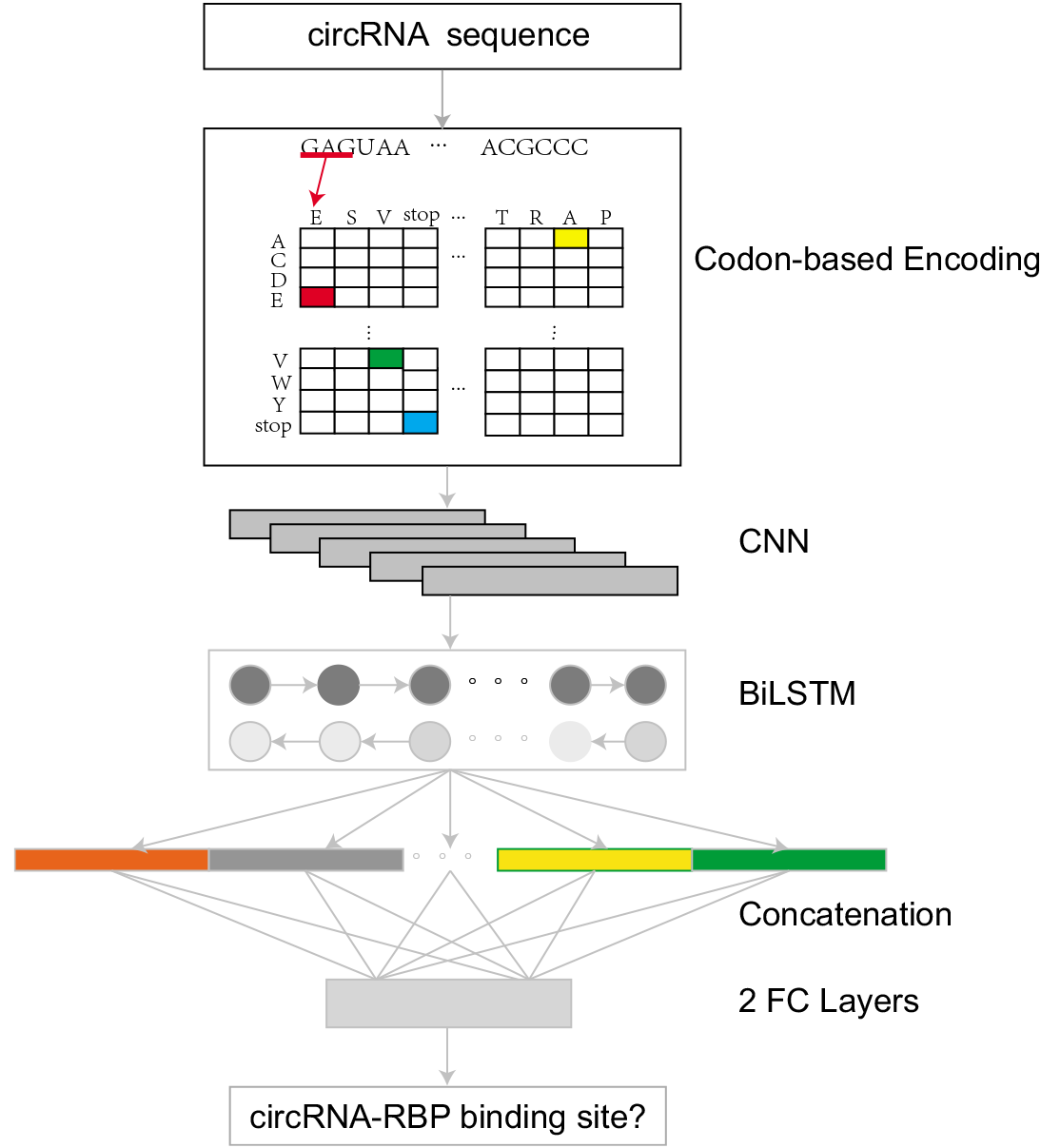
Flowchart of the proposed CRIP method. CRIP represents RNA sequences by stacked codon-based encoding, which is fed into a CNN module followed by a BiLSTM module and further classified through two fully connected layers.

### 2.1 Data preparation

In order to assess the prediction performance of CRIP, we construct a benchmark set of RBP-binding sites on circRNAs. The bound sequences are extracted from the circRNA Interactome database (https://circinteractome.nia.nih.gov/), which houses over 120,000 human circRNAs [11]. Considering that our model conducts prediction based solely on circRNA sequences and a high sequence identity may cause biased result for machine learning methods, we use the CD-HIT package [14] with a threshold of 0.8 to eliminate redundant sequences. Finally, we have a total of 32,216 circRNAs associated with 37 RBPs. For each RBP, we build a classification model, where the positive samples are derived from verified binding sites on circRNAs. Following our previous works [29], we extract sequence segments spanning upstream 50 nt and downstream 50 nt around the binding sites corresponding to the read peaks. Thus each sample is a segment of length 101. The negative samples are extracted from the remaining fragments of the circRNAs, with the same length as positive samples. The positive samples and negative samples are also filtered to remove redundant sequences with a cutoff of 0.8 using CD-HIT. The positive-to-negative ratio is 1:1, and detailed data statistics are listed in Supplementary Table S1.

In addition, since CRIP is also applicable to linear RNA data, we compare the performance of CRIP with the existing tools on the prediction of interactions between linear RNAs and RBPs, using previously published benchmark sets of linear RNAs, i.e., the same dataset used in iONMF [35] and iDeep [29], retrieved from DoRiNA [4] and iCount (http://icount.biolab.si/). There are 31 datasets derived from CLIP-Seq data, corresponding to 31 experiments and covering 19 RBPs. The positive and negative samples are generated in the same way described above, and each of the 31 datasets has 5000 training samples and 1000 test samples, both of which has the positive-to-negative ratio of 1:4.

### 2.2 Stacked codon-based encoding

With the increasing applications of deep learning methods in sequence analysis, traditional feature extraction methods, like k-mer frequency, have been largely replaced by sequence encoding methods. Especially, one-hot encoding has been widely used in biological sequence classification [3]. However, one-hot has obvious drawbacks. For RNA/DNA sequences, each nucleotide is encoded as a 4-dimensional binary vector. Such a low-dimensional feature representation may be incompetent to characterize the sequence information well. Especially, the sequence context information is not encoded in one-hot method.

In order to incorporate context information and get an expanded vector space retaining more sequence features, we propose a new method, called stacked codon-based encoding. Inspired by the recent finding of circRNAs that they can code for proteins [27], we map each 3 consecutive nucleic acids (i.e. 3-mer) in the circRNA sequences into a pseudo amino acid. The mapping is similar to the translation of codons, except that the mapping is conducted in an overlapping manner due to the indeterminacy of the starting site. Also, since this is not a real translation process, we allow stop codons in the middle of the sequence.

Since there are 64 combinations of 3-mers and only 21 different characters (20 amino acid plus a stop codon), each amino acid may correspond to multiple codons. Therefore, our stacked codon-based encoding method can be regarded as a variant of *k*-mer method, where *k* = 3. That is, we extract *k*-mer from the left to right of a sequence using a sliding window in an overlapping way; while unlike *k*-mer method, which leads to a high-dimensional feature space (4*^k^*), we convert the 3-mers into a 21-length alphabet, thus reducing the feature dimensionality and grouping the 3-mers with common biological properties at the same time. Finally, we encode the ‘amino acid’ sequences by the conventional one-hot method, i.e. each amino acid and the stop codon is converted into a 21-D binary vector, where only one feature equals 1.

Let 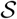 be an RNA sequence of length *L*. It will be converted into a pseudo-amino acid sequence, whose alphabet is 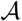=A, C, D, E, F, G, H, I, K, L, M, N, P, Q, R, S, T, V, W, Y, Z, where ‘Z’ denotes the stop codon. 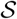 is represented by a 21×(*L*-2)-D matrix **M** (there are a total of *L* – 2 overlapping codons for a sequence of length *L*). The *j*th column of the matrix is a one-hot vector for the *j*th letter in the converted sequence 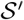, where *j* ∈ {1, 2, · · ·, *L* – 2}. Then the elements of **M** are represented in Eq. (1),

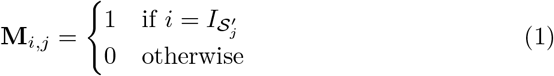

where 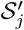 is the *j*th character of 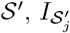 denotes the index of 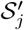 in the alphabet 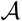 and *i* ∈ {1, 2, · · ·, 21}.

As illustrated in Fig. 1, the corresponding pseudo-amino acid sequence of the input sequence GAGUAA is ESVZ, where GAG codes for E, AGU codes for S, GUA codes for V and UAA is a stop codon. Since the indexes of E, S, V and Z in the alphabet is 4, 16, 18 and 21, respectively, then the generated matrix M is 21×4-D, and **M**_41_, **M**_16,2_, **M**_18,3_ and **M**_21,4_ are equal to 1, while other elements are zero.

For comparison, we consider another coding scheme, called IUPAC [7], which provides another alphabet including 16 characters. IUPAC considers the genetic variation, thus each character in its alphabet corresponds to a polymorphic status of nucleic acids, like ‘A or C’ and ‘not G’, as shown in Supplementary Table S2 [19]. In this paper, the IUPAC method refers to the extended one-hot using the IUPAC alphabet.

### 2.3 The CNN layers

The convolutional neural network (CNN) has been demonstrated to have powerful capability to extract high-level abstract features, not only for image processing but also natural language processing tasks [34]. In this study, we also employ CNN as a feature extractor, whose input is the sequence encoding, i.e., the matrix **M** described in Section 2.2. Assume the size of **M** is *d × l*, where d is the dimensionality of the one-hot vectors (*d* = 21 for the stacked codon-based encoding) and *l* is the length of the converted pseudo-amino acid sequence, we take a one-dimensional convolution along the sequence. For the convolutional layer, we set the length of a filter as *h_f_*, which means that the filters are operated on *h_f_* words/tokens. Let **X***_i_* be the original encoding matrix for the segment of the input sequence processed by the sliding kernels at the *i*th time step, which is actually a submatrix of **M**, consisting of the ith to (*i + h_f_* – 1)-th columns of **M**. For convenience, we take the transposed submatrix of **M** as **X***_i_*. Then the size of **X***_į_* is (*i + h_f_* – 1) × *d*. The corresponding outputs of all **X***_i_*s passing through the *j*-th sliding kernel turns out to be a column vector, *y^j^*. Each element, 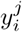 is defined in Eq. (2)

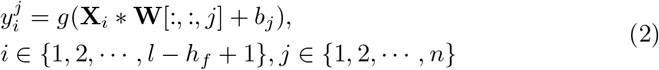

where *g*(·) is a ReLU function, *n* is the number of the filters, **W** is the convolutional filter 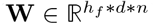, and *b* is the bias. In the pooling layer, we choose an average pooling over the sequence with the length of *h_p_*. The output of the pooling layer for the *j*th filter is a column vector defined as **z***^j^*, where the *m*th element, 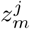, is computed as Eq. (3),

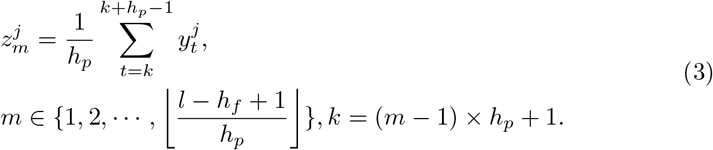

Let **Z** be the matrix whose column vectors are **z***^j^*, i.e., the high-level features learned by the CNN model. **Z** is fed to the subsequent BiLSTM model for classification.

### 2.4 The BiLSTM layer

Through the convolutional filters and average pooling layers, the CNN module learns and integrates local information of RNA sequences. Then we stitch the data of all the channels of each subunit into a new feature vector. In order to further exploit the sequence information, we adopt a bidirectional long and shortterm memory network (BiLSTM). Compared with traditional recurrent neural networks (RNNs), LSTM has advantages in addressing the vanishing/exploding gradient problem and long-term dependency. Especially, BiLSTM exploits the contextual information on both sides. Let s*_t_* and 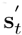 be the hidden states for the forward and backward computation at the *t*th time step. The calculation of s*_t_* and 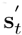 relies on s*_t-1_* and 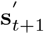, respectively, as defined in Eqs. (4) and (5).

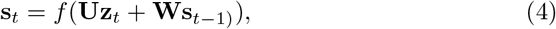

where **U** and **W** are the weight matrices for the input and the hidden state, respectively, and *z_t_* is the input vector in the *t*th step, i.e., the *t*th row vector of *Z*.

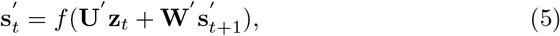

where **U**ʹ and **W**ʹ are the weight matrices for the input and the hidden state used in the backward computation, respectively. To integrate contextual information, the output for *t*th step is defined as,

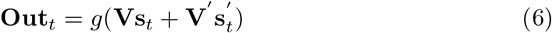

where **V** and **V**ʹ are the transformation matrices of the preceding and following context for the current time step.

### 2.5 Output concatenation and the fully connected layers

By convention, only the last output of LSTM is fed to the fully connected layer for final classification. In this study, we find that the outputs of previous time steps also contain some information helpful for the classification. Therefore, we concatenate the output vectors of all the time points, i.e.,

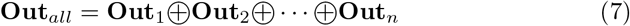

In addition, since the concatenated output has a high dimensionality, we add two fully connected layers to gradually reduce the dimensionality for the final classification, and the softmax layer maps all outputs to probabilities.

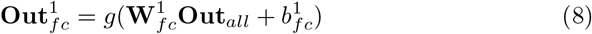

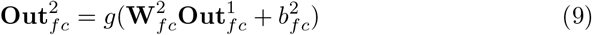

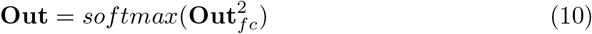

where *g* is the ReLU function, 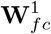 and 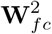 are the weight matrices, and 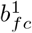 and 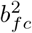 are the bias terms.

### 2.6 Motif analysis

To further explore the sequence patterns of RBP-binding sites on circRNAs, we search motifs from the positive sequence fragments via the MEME suite [2]. The motifs are extracted for each RBP dataset respectively, and the most significant motif corresponding to each RBP is shown in Supplementary Table S3, where the width of motifs ranges from 8 to 15.

## 3 Experimental Results

### 3.1 Experimental settings

In CRIP, the CNN module mainly consists of two parts. The convolution layers have a total of 102 filters with size 7 × 21, and the pooling layers adopt a window with size 5 × 1. The BiLSTM contains 120 neurons. The activation function used in the middle layer is ReLu and in the final output layer is softmax. Adam algorithm is used as the optimizer with learning rate 0.0001, the loss function is categorical cross-entropy, and early stopping is also used to avoid overfitting. The training and test sample ratio is 4:1. And a 5-fold cross-validation is adopted on the training set to select parameters. Through a grid search on the validation set, the batch size of 50 and the training epoch number of 30 achieve the optimum results.

### 3.2 Performance of CRIP

#### Investigation on the feature encoding

In order to represent RNA sequences, both traditional feature extraction methods and sequence encoding schemes have been investigated. For instance, in [33] and [26], *k*-mer frequencies were used as features and classified by SVMs and random forests. Some recent works, DeepBind [1], iDeep [29], and iDeepS [28], adopted one-hot encoding and deep learning models as classifiers.

Here we compare the stacked codon-based encoding with two other sequence encoding methods, i.e. traditional one-hot and IUPAC code [19]. Table 1 shows averaged AUCs of the three encoding methods on the 37 circRNA datasets, working with two different classifiers, namely BiLSTM (i.e., the RNN part in the hybrid neural network) and the hybrid neural network, respectively. In both cases, our method achieves the best performance, and IUPAC outperforms one-hot slightly. Obviously, our method and IUPAC have larger alphabets than the conventional one-hot method, and the extended encoding space is helpful to retain sequence features. As can be seen, benefiting from the CNN module, all the three encoding methods get improved accuracy, and the performance gap becomes smaller compared to using only the LSTM component, suggesting that the deep learning architecture can compensate for the initial simple features.

**Table 1:** Comparison of different encoding methods on 37 circRNA datasets

| Method | Average AUC |  |
| --- | --- | --- |
|  | BiLSTM | Hybrid neural network |
| Codon-based encoding | **0.845** | **0.872** |
| One-hot | 0.821 | 0.862 |
| IUPAC | 0.828 | 0.868 |

#### Investigation on the learning models

As described in Sections 2.3 and 2.4, our model includes a CNN module and a BiLSTM module. In order to evaluate the contribution of each module in the hybrid neural network, we examine the performance of the single modules, respectively. The results are shown in Fig. 2.

**Fig. 2:**
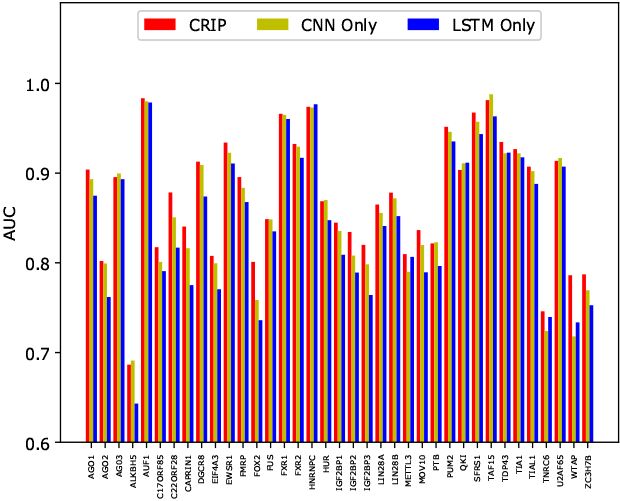
The AUCs of the LSTM module, CNN module and the hybrid model on 37 circRNA datasets.

Apparently, the single modules do not perform as well as the hybrid model which has an average AUC of 0.872. The CNN module is better than the BiLSTM module (0.861 vs. 0.845). It shows that CNN does have an advantage in feature extraction and can extract more accurate information for the detection of RNA-protein interaction.

### 3.3 Comparison with the existing predictors for predicting circRNA-RBP interactions

#### Comparison with traditional machine learning methods

Regardless of space structure, the identification of RBP-binding sites on RNAs relies on the same information source, namely RNA sequences, for any types of RNAs. Thus, we apply previous methods designed for linear RNAs to the circRNA data. Especially, in [26], the authors proposed RPISeq-SVM and RPISeq-RF to identify RNA-protein interactions, which used two representative shallow learning models, SVMs and random forests. And the RNA sequence features were represented by normalized 4-mer composition. Here we implement these two methods, which are trained and evaluated using the same 37 circRNA datasets as CRIP. The results are shown in Fig. 3.

**Fig. 3:**
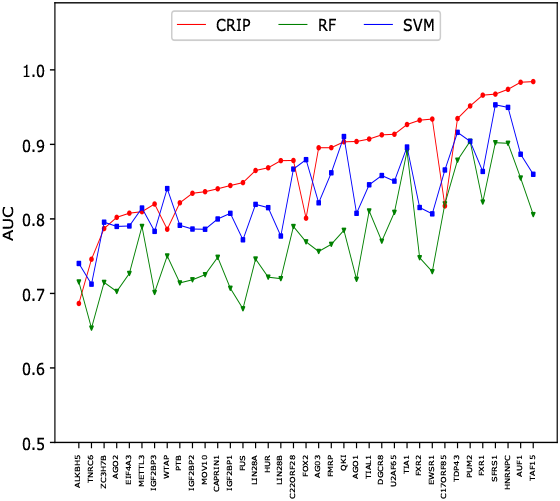
Comparison of AUCs between CRIP and the predictors based on traditional machine learning models on 37 circRNA datasets.

The advantages of CRIP over the traditional learning methods are obvious. Among the 37 datasets, CRIP achieves the best results on 30 datasets. The average AUC of CRIP is 0.872, which is 4.6% higher than that of the SVM (0.834), and 13.4% higher than that of RF (0.769), demonstrating the advantages of the proposed deep model against traditional learning methods.

#### Comparison with the existing deep learning methods

To further assess the performance of CRIP, we compare it with the deep learning based predictors using sequences alone, including DeepBind [1] and iDeepS [30], while iDeep and iDeepE are excluded in the comparison due to the following reasons: i) iDeep requires annotation information of gene regions and clip-cobinding [29]; ii) iDeepE breaks sequences into multiple overlapping subsequences with size 101, which yields better performance than DeepBind for long sequences but performs similarly to DeepBind for short sequence fragments. DeepBind utilizes only sequence features and adopts a sequence CNN, while iDeepS integrates both sequence and secondary structure information, and adopts a similar model architecture as CRIP. Since these two methods were designed for linear RNAs rather than circRNA, to perform a fair comparison, we conduct the experiments on the benchmark sets for linear RNAs, including 31 datasets. The results are shown in Fig. 4.

**Fig. 4:**
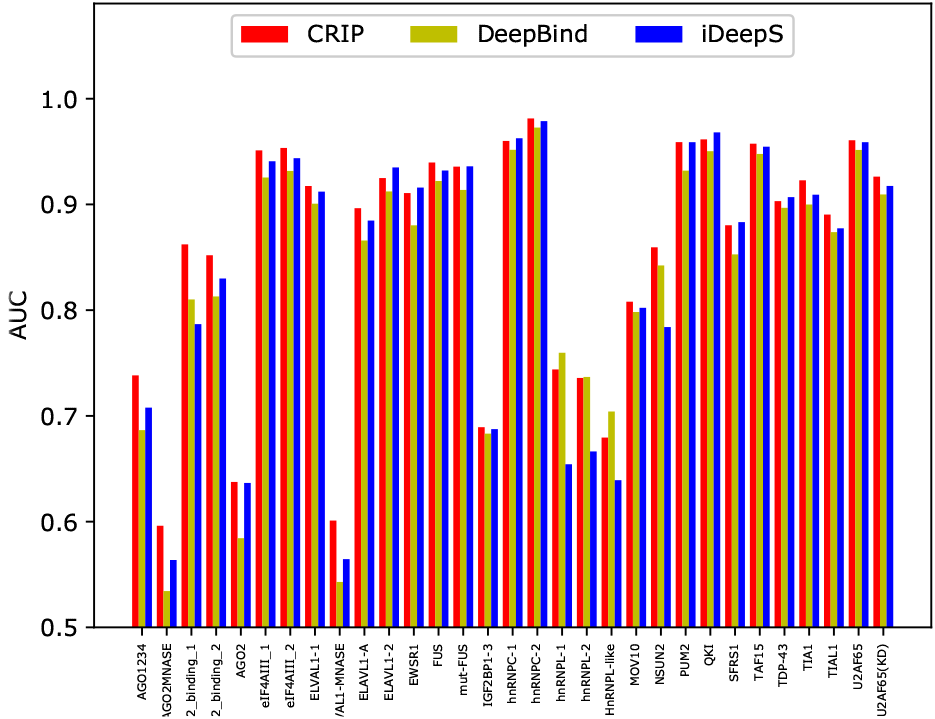
Comparison of the AUCs of DeepBind, iDeepS and CRIP on 31 linear RNA datasets

Apparently, CRIP has achieved the best results for most of the RBPs. For some RBPs, like the Argonaute family of proteins (AGO), CRIP has a much higher AUC than the two other methods. iDeepS has a slight advantage over DeepBind, perhaps because the secondary structure information incorporated in iDeepS helps improve the prediction accuracy. On average, CRIP obtains an AUC value of 0.856, which is higher than that of DeepBind (0.835) and iDeepS (0.839). This experiment demonstrates that CRIP not only applies to linear RNAs, but also improves AUC for about 2% compared with the state-of-the-art deep models designed for linear RNAs.

### 3.4 Experiments for the RBPs common to circRNAs and linear RNAs

#### The models trained on linear RNAs could not simply be applied to circRNAs

Previous studies have employed both shallow and deep learning models for predicting RNA-protein interactions, but none of them was designed for circRNAs. Note that there are some RBPs shared by circRNAs and other types of RNAs, thus we compare the RBPs used in this study and previous studies [28][30] and find 11 RBPs common to linear RNAs and circRNAs. We first check whether the predictors trained by linear RNAs can be generalized to circRNAs.

For each circRNA test set of the 11 shared RBPs, we compare the performance of CRIP with iDeep*^1^. Fig. 5 shows the prediction results of iDeep* and CRIP for the common RBPs. As can be seen, CRIP outperforms iDeep* on all of the 11 datasets. Especially for FUS, HNRNPC and MOV10, CRIP improves the AUC by more than 10%, indicating that the training sequences in iDeep^*^ (linear RNAs) may be very different from the test sequences (circRNAs), and these two types of RNAs may differ in the interaction mechanisms to the same RBPs.

**Fig. 5:**
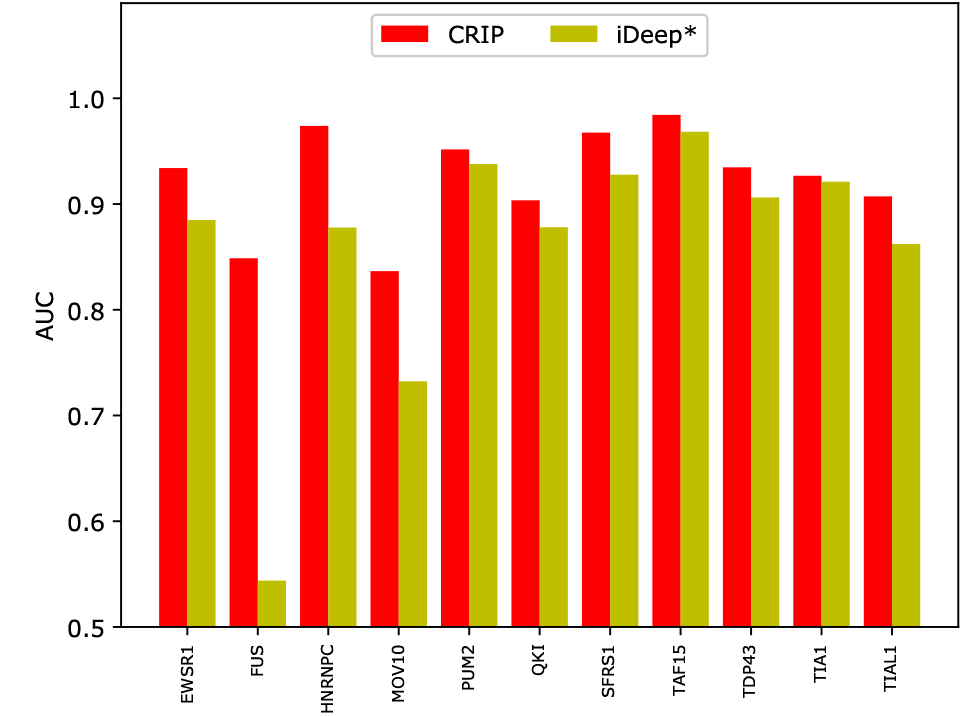
Comparison of AUCs of CRIP and iDeep on the common RBPs. iDeep* is trained on linear RNAs and evaluated on test circRNAs. CRIP is trained on training circRNAs and evaluated on test circRNAs.

This experiment demonstrates the necessity of developing a specific predictor for identifying binding sites in circRNA sequences, not only for new RBPs, but also for the shared RBPs with linear RNAs.

#### Linear RNAs and circRNAs binding to the same RBPs differ in prediction accuracy

Compared with linear RNAs, circular RNAs have their distinct structure and mechanism for binding to proteins, which motivates us to develop a new tool for identifying circRNA-protein interactions, while CRIP can actually be applied to any type of RNAs. As mentioned in Section 3.4, circRNAs share some RBPs with linear RNAs. Therefore, we compare the prediction performance of CRIP on the common RBPs for linear RNAs and circRNAs, as shown in Fig. 6.

**Fig. 6:**
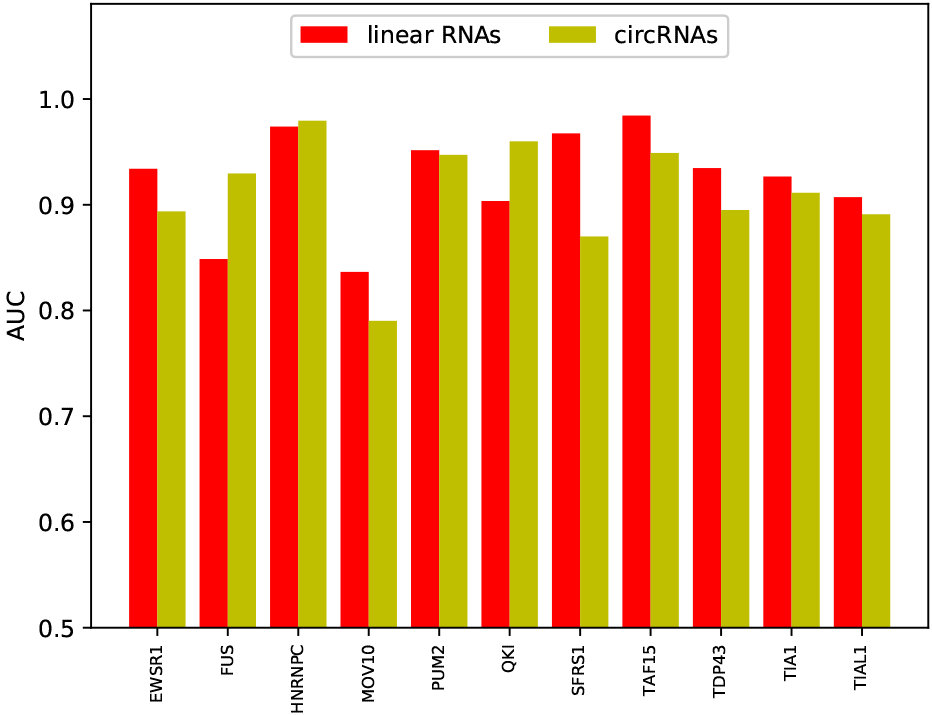
Comparison of AUCs for common RBPs shared by linear RNAs and circRNAs, where the AUCs for linear RNAs and circRNAs are obtained both by CRIP using sequence information.

Among the 11 common RBPs, circRNAs have a lower accuracy on 8 datasets. The performance of linear RNAs and circRNAs differs a lot for some RBPs, e.g. FUS and SFRS1. The reasons for the performance difference are manifold. As a machine learning-based method, the performance of CRIP heavily relies on the scale of training data. In the linear RNA dataset, each RBP corresponds to 10000 sequence segments in the training set (5000 positives and 5000 negatives); while when we construct the circRNA dataset, we extract fragments from all binding sites, thus if there is a large number of binding sites on a circRNA, then it will have abundant training samples; otherwise the training set will be small. In other words, our datasets vary in scale. Generally, a large training set will lead to good performance. For instance, the FUS and HNRNPC datasets of circRNA binding sites are the two biggest ones among the 11 RBPs, with 20000 and 14224 positive samples, respectively. Both of them achieve better accuracy compared against their corresponding datasets of linear RNAs. Especially, for FUS, the AUCs of circRNA data and linear RNA data are 0.930 vs. 0.849. However, there are a few exceptions. On the QKI dataset, CRIP also performs better on circRNAs than on linear RNAs (0.960 vs. 0.904), but the number of positive training samples is only 1033, suggesting that there are other factors affecting the prediction accuracy. Through a motif search using the MEME suite, we identify a conserved and concentrated pattern, ‘ACUAAC’, on the circRNAs binding to QKI. This motif has been verified and was included in the CISBP-RNA database[32]. Similarly, we also find two other conserved fragments, which are consistent with the motifs in CISBP-RNA, namely ‘UGUA’ for the binding to Pum2 and ‘UUUU’ for the binding to TIA1 (Table 2). These two datasets have only 2829 and 2202 positive samples, respectively, while their accuracies are very close to those of their corresponding sets on linear RNAs. From these observations, we conclude that conserved motifs may also help in the performance enhancement.

**Table 2:**
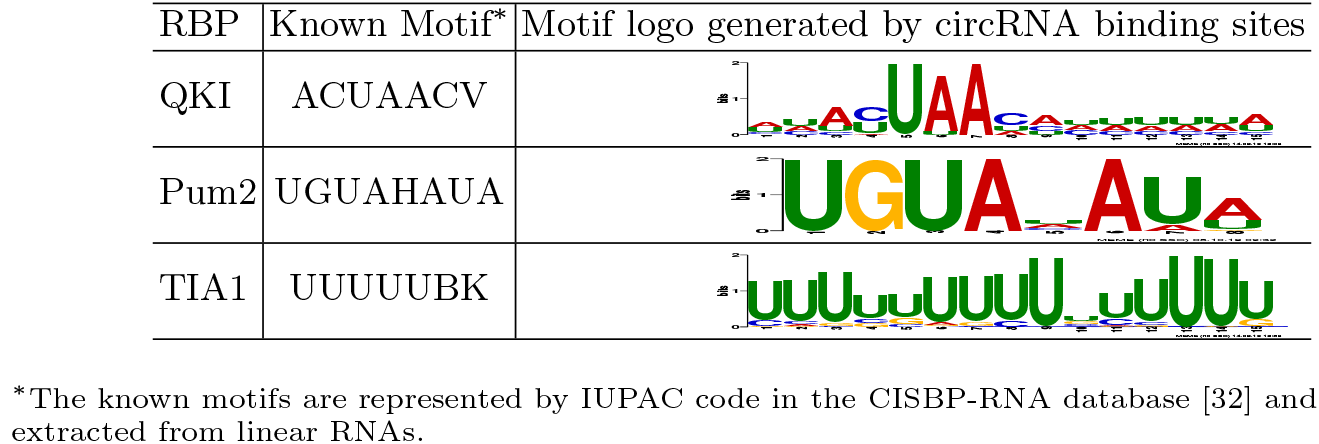
Some common motifs shared by circRNAs and linear RNAs.

## 4. Discussion

This study explores the potential of predicting circRNA-RBP interactions based on RNA sequences using a deep neural network model. The main contributions include:

- Construct benchmark datasets of circRNA segments binding to RBPs, and propose the first specific predictor for the identification of RBP binding sites on circRNAs;
- Design a new encoding scheme to represent RNA sequences, and successfully apply it to the prediction of RBP binding sites on both circRNAs and linear RNAs;
- Employ a hybrid deep neural network to further enhance the performance.

The success of CRIP lies in the enriched feature representation and powerful DNN model. Unlike the conventional one-hot method which encodes nucleic acids one by one, the new encoding method traverses the 3-mers sequentially in an overlapping manner, just like the traditional *k*-mer feature extraction, and then encode the 3-mers into a binary vector according to the codons for amino acids. Thus, benefiting from the context information retained in the 3-mers and the expanded feature space, the new stacked-codon encoding provides more informative representations. Moreover, the CNN and BiLSTM components further learn high-level abstract features and contextual information from the encoding vectors, respectively.

Besides the evaluation of the new model, we perform motif analysis for circRNAs interacting with 37 different RBPs. Specifically, we use the MEME tool [2] to extract conserved patterns from the binding sites. Supplementary Table S3 shows the most significant motifs (according to E-value) extracted from the positive samples for each RBP. As mentioned in Section 3.4, we find that some circRNAs have the same motifs as linear RNAs when binding to the same RBPs, indicating the two types of RNAs may share a common binding mechanism. We also find that the binding sites for some RBPs exhibit a common pattern, i.e. “GAAGAAG”, including AGO2, ALKBH5, CAPRIN1, LIN28B, IGF2BP3, etc. Actually, it is a common motif related to RNA modification [22, 8].

Despite the common motifs, for the same RBP, binding sites on circRNAs and linear RNAs may have large sequence diversity. A typical example is the dataset for binding to FUS. The classifier trained on linear RNAs has a very low prediction accuracy on circRNAs (Fig. 5); while using CRIP which is trained on circRNAs, the accuracy becomes much higher than that of linear RNAs. Therefore, it would be interesting to explore the different binding mechanisms that lead to performance variance.

In addition, there exist some potential applications of CRIP. RBPs have been discovered to play important roles in circRNA production. To identify those RBPs related to circNRA biogenesis, we need to first predict the interactions between circRNAs and all available RBPs. As some RBPs, e.g. FUS [12] and QKI [6], have been experimentally verified to be involved in circRNA biogenesis, through the analysis on the binding sequence patterns from these verified interactions, we can identify novel RBPs involved in circRNA biogenesis.

## 5 Conclusions

This study aims to identify circRNA-protein interaction sites by using a machine learning model. By treating the task as a binary classification problem, we propose a new sequence encoding scheme and a hybrid neural network model. The idea of the new encoding method is to convert RNA triplets into pseudo-amino acids based on nucleotide codons in an overlapping manner, and represent the pseudo-amino acids via one-hot encoding. And the hybrid neural network consists of a CNN module and a BiLSTM module. The goal of using a hybrid model is to combine the advantages of both the two deep architectures, and obtain better high-level abstraction feature representations for the classification. The experiments show that both the new sequence encoding method and the hybrid model contribute to the performance improvement. Compared to the existing predictors, our model has an obvious advantage in the prediction accuracy. We believe that this tool will contribute to uncovering functions of circRNAs.

## Footnotes

1 Since circRNAs only have sequence information, we retrain the iDeep [29] using linear RNA sequences alone and name it iDeep*.

