## Supplementary Table S1, Supplementary Table S2, Supplementary Table S3 for "Predicting circRNA-RBP interaction sites using a codon-based encoding and hybrid deep neural networks"

### Supplementary Materials

#### 1 Tables

Table S1. Data statistics of the datasets

| RBP | Positive # | Negative # |
|---|---|---|
| AGO1 | 17318 | 17318 |
| AGO2 | 20000 | 20000 |
| AGO3 | 3124 | 3124 |
| ALKBH5 | 770 | 770 |
| AUF1 | 2896 | 2896 |
| C17ORF85 | 1016 | 1016 |
| C22ORF28 | 5292 | 5292 |
| CAPRIN1 | 5298 | 5298 |
| DGCR8 | 20000 | 20000 |
| EIF4A3 | 20000 | 20000 |
| EWSR1 | 4695 | 4695 |
| FMRP | 20000 | 20000 |
| FOX2 | 605 | 605 |
| FUS | 20000 | 20000 |
| FXR1 | 927 | 927 |
| FXR2 | 5635 | 5635 |
| HNRNPC | 14224 | 14224 |
| HUR | 20000 | 20000 |
| IGF2BP1 | 20000 | 20000 |
| IGF2BP2 | 10000 | 10000 |
| IGF2BP3 | 20000 | 20000 |
| LIN28A | 18277 | 18277 |
| LIN28B | 7888 | 78888 |
| METTL3 | 2666 | 2666 |
| MOV10 | 5889 | 5889 |
| PTB | 20000 | 20000 |
| PUM2 | 2829 | 2829 |
| QKI | 1033 | 1033 |
| SFRS1 | 8595 | 8595 |
| TAF15 | 1467 | 1467 |
| TDP43 | 5484 | 5484 |
| TIA1 | 2202 | 2202 |
| TIAL1 | 5456 | 5456 |
| TNRC6 | 1101 | 1101 |
| U2AF65 | 8224 | 8224 |
| WTAP | 496 | 496 |
| ZC3H7B | 13119 | 13119 |

Table S2. IUPAC codes for nucleotide bases

| Symbol | Mnemonic | Translation |
|---|---|---|
| A | | A (adenine) |
| C | | C (cytosine) |
| G | | G (guanine) |
| T | | T (thymine) |
| U | | U (uracil) |
| R | puRine | A or G (purines) |
| Y | pYrimidine | C or T/U (pyrimidines) |
| M | aMino group | A or C |
| K | Keto group | G or T/U |
| S | Strong interaction | C or G |
| W | Weak interaction | A or T/U |
| H | not G | A, C or T/U |
| B | not A | C, G or T/U |
| V | not T/U | A, C or G |
| D | not C | A, G or T/U |
| N | any | A, C, G or T/U |

Table S3. The corresponding binding motifs in RNA sequences for the 37 RBPs

| | | | | | |
|---|---|---|---|---|---|
| AGO1 | 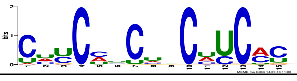 | AGO2 | 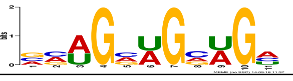 | AGO3 | 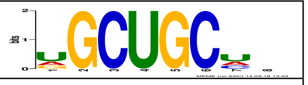 |
| ALKBH5 | 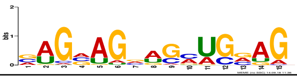 | AUF1 | 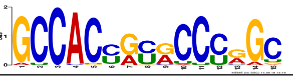 | C17ORF85 | 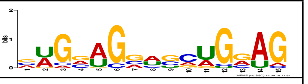 |
| C22ORF28 | 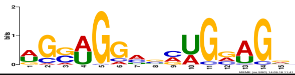 | CAPRIN1 | 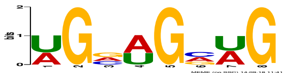 | DGCR8 | 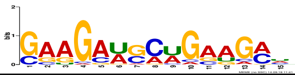 |
| EIF4A3 | 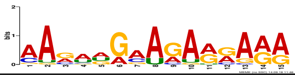 | EWSR1 | 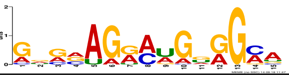 | FMRP | 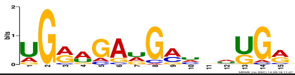 |
| FOX2 | 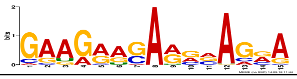 | FUS | 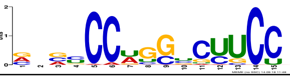 | FXR1 | 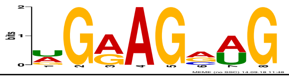 |
| FXR2 | 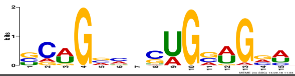 | HNRNPC | 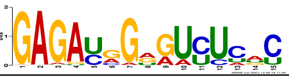 | HUR | 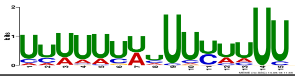 |
| IGF2BP1 | 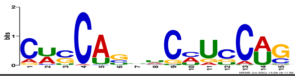 | IGF2BP2 | 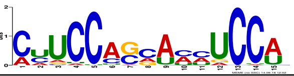 | IGF2BP3 | 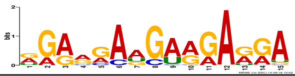 |
| LIN28A | 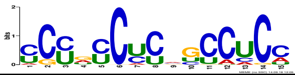 | LIN28B | 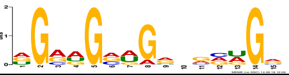 | METTL3 | 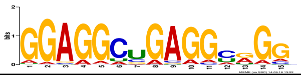 |
| MOV10 | 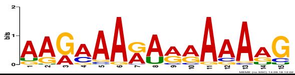 | PTB | 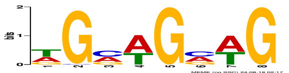 | PUM2 | 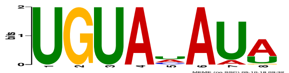 |
| QKI | 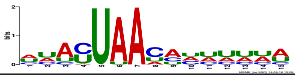 | SFRS1 | 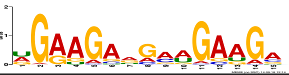 | TAF15 | 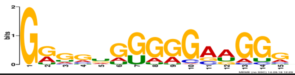 |
| TDP43 | 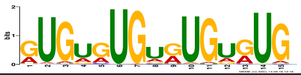 | TIA1 | 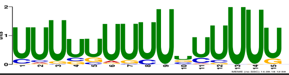 | TIAL1 | 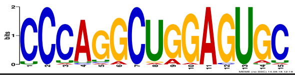 |
| TNRC6 | 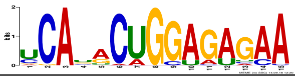 | U2AF65 | 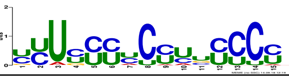 | WTAP | 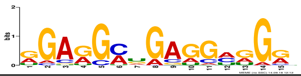 |
| ZC3H7B | 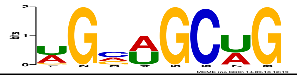 | | | | |
